# Host-mediated pH influences microbiome assembly and function on the phylloplane

**DOI:** 10.64898/2026.03.21.713394

**Authors:** Jean-Baptiste Floc’h, Cristal Lopez-Gonzalez, Tanya Renner, Kadeem J. Gilbert

**Affiliations:** Department of Plant Biology, W. K. Kellogg Biological Station, Program in Ecology, Evolution, & Behavior, and Plant Resilience Institute, Michigan State University, Hickory Corners, MI, USA; Faculty of Biological and Environmental Sciences, University of Helsinki, Helsinki, Finland; Department of Entomology, The Pennsylvania State University, State College, PA, USA

**Keywords:** Phylloplane, phyllosphere, pH, metatranscriptomics, *Gossypium*, *Beta vulgaris*, *Nepenthes*

## Abstract

Plant leaves harbor diverse microbial communities influenced by environmental inputs and host traits, yet it remains unclear whether leaves act as passive substrates or active ecological filters that reorganize microbial functional capacity. Phylloplane pH regulation is one hostplant trait that has been traditionally underexplored. We used metatranscriptomics to examine microbial gene expression on the phylloplane and within whole leaves of five plant species spanning the extremes of baseline phylloplane pH, including hyperalkalinizing *Gossypium* species, weakly buffering *Beta vulgaris*, and hyperacidifying *Nepenthes* species. Young leaves were inoculated with a common soil-derived microbial community to quantify host-associated restructuring of taxonomic and functional profiles, and short-term pH perturbations were applied to test the effect of transient abiotic stress. Across both phylloplane and whole-leaf datasets, host species identity was the primary axis structuring microbial taxonomic composition and expressed functional repertoires. Leaf-associated communities diverged from the source inoculum, but retained a substantial shared functional backbone enriched for central biosynthetic and core metabolic pathways. Host-associated differentiation reflected selective retention and redistribution of reactions within this shared environmental pool rather than acquisition of novel metabolic capacity. Enriched pathway subsets were metabolically coherent and taxonomically distributed across multiple bacterial orders, consistent with functional redundancy and trait-based assembly. Among hosts, *Gossypium* exhibited the strongest restructuring relative to inoculum, suggesting comparatively stronger host-associated filtering. In contrast, short-term pH manipulation did not induce consistent community-wide functional reorganization. Microbial physiological responses to the phylloplane environment and external pH were observed at the organismal level. Together, these results position leaves as active ecological filters that reorganize microbial functional landscapes through host-specific trait regimes. This work begins to implicate some role of phylloplane pH regulation in microbial assembly and function.

## Introduction

Microbial communities associated with plants emerge from the interplay between environmental conditions and host traits. In belowground systems, particularly the rhizosphere, host genotype and physiology are recognized as dominant forces shaping microbial assembly and function [1,2]. By contrast, aboveground leaf surfaces are often conceptualized primarily as habitats structured by external abiotic drivers such as temperature, ultraviolet radiation, hydration dynamics, and atmospheric deposition [3–5]. Whether leaves function mainly as passive substrates exposed to fluctuating environmental inputs or instead act as active ecological filters that reorganize microbial functional capacity (i.e., the repertoire of functions encoded and potentially expressed by the microbial community) is still understudied.

Among abiotic parameters influencing microbial systems, pH is one of the most powerful determinants of microbial growth, physiology, and community structure across soils, aquatic systems, and animal-associated microbiomes [6–9]. Critically, pH is not merely an external environmental variable; it can be modified and actively regulated by host organisms [10,11] and can thus be understood as an extended host phenotypic trait as well. In the rhizosphere, proton extrusion and ion exchange alter local pH, influencing nutrient availability and microbial assembly [12,13]. While underappreciated, leaves can also rapidly alter local pH by transporting protons and cations to the leaf surface (known as the phylloplane), generating species-specific phylloplane pH regimes, though most plants are typically around neutral [14,15]. In carnivorous plant species, acidic environments are well documented (reaching as low as pH <2.0), whereas members of the Malvaceae can alkalinize the phylloplane to extreme levels (up to pH >10), thus baseline phylloplane pH spans extremes across angiosperm diversity [16–18]. These striking interspecific differences suggest that phylloplane pH regulation may represent an underexplored axis of host-mediated microbiome structuring. Further, environmental pH is dynamic; even short-term changes can be fatal for microbes if exposed to pH levels outside of their physiological limits. Only 10 minutes of direct exposure to pH 2 can kill *Helicobacter pylori* colonies [19]. Sublethal changes in the pH environment may also hypothetically alter the functional profile of a microbial community, as the expression of many molecular processes (e.g., ion transport) can depend on external pH gradients. Thus, the ability of some plants to rapidly buffer against external pH changes [17] may override environmental perturbation in determining the survival and persistence of microbes colonizing the phylloplane.

Despite increasing recognition that host identity influences phyllosphere community composition [5,20], most work has focused on taxonomic patterns rather than functional organization. Yet microbial ecosystems frequently display functional redundancy, wherein taxonomically distinct communities maintain overlapping metabolic capabilities [21,22]. From a trait-based ecological perspective [23] of microbes, environmental filtering [24] may therefore operate not by generating entirely novel metabolic repertoires, but by reshaping the distribution and expression of functions within a shared environmental pool. In the phyllosphere, characterized by nutrient limitation, oxidative stress, desiccation, and microscale chemical heterogeneity [3,4], understanding host influence requires moving beyond species richness toward functional architecture and physiological responses.

Metatranscriptomics provides an avenue to evaluate this functional architecture by capturing community-level gene expression profiles under specific host and environmental contexts [25,26]. Furthermore, it also allows for the opportunity to link microbial gene expression to the taxonomic composition of their communities and to infer physiological processes [27,28]. This functional resolution enables explicit testing of alternative models of host control over microbial communities [29,30]. Specifically, reaction- and pathway-level analyses allow discrimination between two contrasting models of host influence: 1) hosts assemble distinct metabolic repertoires unique to each species, or 2) hosts selectively reorganize a largely shared environmental functional backbone. Distinguishing between these alternatives is essential for understanding how plant traits shape microbiome function at ecosystem-relevant scales.

Beyond resolving functional structure, metatranscriptomics also provides a means to investigate microbial physiological responses to environmental perturbations within short timescales [31,32]. Because transcriptional activity responds rapidly to abiotic change, this approach offers organism-level insight into stress responses and functional plasticity that are often obscured in community-level analyses [33,34].

Here, we used metatranscriptomics to test whether leaf-associated microbial communities are structured primarily by host identity and whether interspecific differences in phylloplane pH regulation correspond to differences in functional organization. We examined five plant species representing a range of phylloplane regulation pH strategies, spanning extremes: hyperalkalinizing *Gossypium arboreum* and *G. hirsutum*, weakly buffering *Beta vulgaris*, and hyperacidifying *Nepenthes bicalcarata* and *N. rafflesiana*. To isolate host-associated effects, young leaves were inoculated with a common soil-derived microbial community. We further applied short-duration pH perturbations to evaluate whether transient abiotic shifts override host-structured organization, or whether the ability of certain hostplants to buffer phylloplane pH obviates this external influence. We also aimed to observe whether any functional shifts can be directly linked to microbial physiological responses to pH. It is important to note that the goal of our phylloplane epimicrobiome experiment was not to mimic natural microbial colonization processes or realistic acid rain events, but rather to assess the potential of host filtering and phylloplane pH regulation at extremes. Microbial transcripts were analyzed from these experimental epimicrobiome communities, as well as from independent whole-leaf datasets, to compare functional structuring across leaf tissues, and the potential for inoculum-derived microbial functions to converge upon functions present in uninoculated leaves. By integrating taxonomic composition, reaction-level repertoires, pathway enrichment, and taxonomic contribution analyses, we assess whether leaves construct distinct metabolic communities or instead reorganize a shared environmental functional reservoir. This framework clarifies the relative roles of host identity and short-term pH perturbation in shaping microbial functional landscapes in the phyllosphere.

## Methods

### Epimicrobiome experiment

#### Plant material and growth conditions

We selected five hostplant species spanning a broad range of phylloplane pH regulation capacities [14]: *Gossypium arboreum* (PI 615701) and *G. hirsutum* (PI 529181), known to exhibit hyper-alkalinizing leaf surfaces; *Beta vulgaris* (Ames 3060) with near-neutral surface pH; and the carnivorous species *N. bicalcarata* (BE-3031) and *Nepenthes rafflesiana* (BE-3722), known to exhibit hyper-acidifying leaf surfaces.

Seeds of *Gossypium* and *Beta* were obtained from the USDA Germplasm Resources Information Network (GRIN), and *Nepenthes* plants were derived from micro-propagated clones (Carnivero, Austin TX; source: Borneo Exotics Ltd.). *Gossypium* and *Beta* were grown in a greenhouse (16 h light/8 h dark; 24 °C day / 21 °C night). *Nepenthes* were grown in a Conviron PGR15 growth chamber (12 h light/12 h dark; 28 °C day / 26 °C night; 80% relative humidity; 250 µmol m⁻² s⁻¹). Plants were grown in a standardized peat-vermiculite or sphagnum (for *Nepenthes*) substrate and watered with a shared source of distilled deionized water (ddH_2_O). Experiments were conducted on the youngest fully expanded leaves (2-3 weeks post-germination) to minimize ontogenetic variation. Leaves were not surface sterilized, in order to avoid disrupting normal phylloplane physiology; however, RNAseq trials using uninoculated leaves yielded no detectable microbial RNAs from the phylloplane.

#### Soil slurry inoculation and pH manipulation experiment

To establish a common microbial inoculum across host species, we prepared a soil slurry from a silty clay loam soil collected from a wooded area on The Pennsylvania State University campus (40°48’4.06”N, 77°52’1.72”W). Soil (pH ∼6.87) that was mixed 1:1 with molecular-grade water, homogenized, and filtered through a 0.45 µm vacuum filter to remove large particles while retaining microbial cells.

Leaves were immersed in sterile, nuclease-free microcentrifuge tubes containing the slurry for three days to allow colonization (sealed with parafilm to prevent aerial colonization). Each leaf represented one replicate community (n = 6 per host, except *N. bicalcarata* n = 4). Control, parafilm-sealed slurry tubes not in contact with leaves were placed near the pots and maintained in parallel to monitor potential stochastic community changes through time.

Following the inoculation period, leaves were treated with either pH 6.5 or pH 2.0 solutions (solutions of ddH_2_O and HCl, vacuum filtered at 0.22 μm to exclude contaminating cells) applied by sterile swabbing (Texwipe® TX®761 Alpha® series) of both the adaxial and abaxial surfaces, and then these treated leaves were collected for microbial collection after 5 minutes of pH treatment exposure, removed from the plant using sterile (autoclaved, UV-crosslinked, and wiped with RNAse-Away) forceps. To measure baseline phylloplane pH, we measured the pH of the remaining slurry from the inoculum tubes removed from the leaves, using a flat-tipped pH probe (HI981037; Hanna Instruments Inc.); baseline pH was measured on the same day as leaf surface microbe collection.

#### RNA extraction and sequencing for epimicrobiomes

We collected microbial cells by sonicating (Bronson® Ultrasonics, Emerson Electric) leaves in tubes of 200 μL DNA/RNA Shield™ (Zymo Research) for 20 minutes, incubated the preserved cell samples at 4°C for 24 hours for maximal cell wall penetrance, and then extracted RNA using the ZymoBIOMICS DNA/RNA Microprep Kit (Zymo Research), without mechanical lysis (to minimize RNA degradation), and an added Proteinase K treatment to maximize nucleotide recovery. Additionally, we added an extra drying step after the final wash step in the kit protocol and incubated the 10 μL nuclease-free H_2_O on the filter for 1 minute before elution. RNA was quantified using a Qubit 1.0 Fluorometer and a Qubit High Sensitivity (HS) Assay Kit (Thermo Fisher Scientific). RNA quality and concentration were measured using a NanoDrop microvolume spectrophotometer (Thermo Fisher Scientific). RNA extractions were sent to The Penn State College of Medicine Genome Sciences Core (Hershey, PA, USA), where library preparation and sequencing (100 bp paired-end) was performed on an Illumina NovaSeq 6000 platform (Illumina, Inc.).

### Whole-leaf microbiome dataset

For comparison, we analyzed previously generated whole-leaf transcriptomes from an independent pH exposure experiment using the same hostplant species, same accessions, and clones [17]. Briefly, the set of hostplants leaves (*G. arboreum*, *G. hirsutum*, *B. vulgaris*, *N. bicalcarata*, *N. rafflesiana*) were sprayed with pH 6.5, 4.0, or 2.0 solutions for 5 minutes, or left untreated in case of dry controls. Total mRNA was extracted and sequenced (for more details see [17]). Raw reads (GEO accession GSE281272) followed the same bioinformatic analysis described below as the metatranscriptome of the epimicrobiome experiment, to extract the microbial reads portion (from “contaminants” removed from plant transcriptome assembly). These transcriptomes were not specifically designed for microbial profiling; microbial reads detected in these datasets were interpreted as the potential functional profile being derived from resident endophytic or surface-associated communities.

### Bioinformatics

#### Quality control and hostplant read removal

Reads were processed using FastQC v0.12.1, MultiQC v1.4, and Cutadapt v4.4 (Phred ≥ 20). Hostplant reads were removed using Kneaddata v0.12.0, using a custom multispecies reference database created with available hostplant reference genomes using nucleotide BLAST (blastn) [35]: *G. arboreum* [36], *G. hirsutum* [37], *B. vulgaris* [38], *N. gracilis* [39]. rRNA sequences were removed using SortMeRNA v4.3.6 (SILVA databases v138) [40].

#### Taxonomic profiling

Taxonomic composition was inferred using MetaPhlAn4 (mpa_vJun23_CHOCOPhlAnSGB_202403 database) [41]. Relative abundances were computed using default marker-based normalization. Species with <0.01 relative abundance were excluded from visualization with Graphlan v1.1.3.

#### Functional profiling

Functional profiling was performed using the HMP Unified Metabolic Analysis Network (HUMAnN) v3.0 [42] against UniRef50 and MetaCyc databases (May 2024 release). Gene families were regrouped to: MetaCyc reactions (RXNs), MetaCyc pathways and Gene Ontology (GO) terms. Abundances of RXNs, pathways, and GO terms were normalized to counts per million (CPM). Features with <0.01% abundance were excluded prior to statistical analyses.

### Statistical Analyses

All analyses were conducted using R Statistical Software v4.2.3 [43]. Alpha diversity (Shannon and Inverse Simpson) and Bray-Curtis dissimilarity were calculated using vegan v2.6-4 [44]. Differences were tested using Kruskal-Wallis tests or ANOVA as appropriate.

Community structure was evaluated using redundancy analysis (RDA) on Hellinger-transformed abundance matrices (decostand(method = “hellinger”)). Models included: taxonomic profiles: (tax_mat ∼ hostplant + treatment + hostplant:treatment); reaction profiles (rxn_mat ∼ hostplant + treatment + hostplant:treatment). Significance was assessed using 999 permutations (anova.cca). Partial RDAs testing pH effects while conditioning on host were specified as: rxn_mat ∼ baseline_pH + Condition(hostplant). Permutation schemes were restricted within host species where appropriate. Distance-to-inoculum was estimated by calculating Bray-Curtis distances between each leaf sample and inoculum replicates. Differences among hosts were tested using Kruskal-Wallis and ANOVA.

#### Reaction repertoire intersection analysis

To evaluate restructuring of the functional repertoire, reaction-level CPM tables were converted to group-level presence-absence matrices. Reactions were called present within a sample if CPM ≥ 1. For each host/treatment, reaction prevalence was calculated as the number of samples with presence (k) out of total samples (n). In the primary analysis, we applied a permissive detection-based screen (k ≥ 1; CPM ≥ 1) to retain reactions detected in at least one replicate per group. As a stricter sensitivity analysis, we required reactions to be present in ≥66% of samples within a group (k/n ≥ 0.66) and in at least two samples (k ≥ 2; CPM ≥ 1). Group-level binary presence-absence matrices were visualized using UpSetR/ComplexHeatmap intersection plots, and repertoire turnover relative to inoculum was quantified using Jaccard distance on the resulting presence-absence profiles.

#### Pathway enrichment analysis

Reactions were mapped to MetaCyc pathways using reaction-to-pathway annotations from Pathway Tools v29.5 [45]. For intersections containing ≥10 reactions, enrichment was tested using a one-sided hypergeometric test with background defined as the union of all annotated reactions across tested intersections. P-values were corrected using Benjamini-Hochberg FDR (q ≤ 0.05).

### Differential abundance testing (MaAsLin3)

Pathway and GO-term abundances were analyzed using a microbiome multivariable associations with linear models tool (MaAsLin3) v1.10.0 [46]. The model formula included hostplant species and pH treatment (feature ∼ host + treatment; normalization: TSS; transformation: log; method: linear model). Significance threshold: FDR < 0.05 (Benjamini-Hochberg). Host × treatment interactions were tested separately. Output tables from MaAsLin3 were used to identify differentially abundant pathways or GO terms linked to each host per treatment compared to inoculum.

#### Taxonomic contribution analysis

HUMAnN3 stratified reaction outputs were aggregated at taxonomic order level. For each pathway, reaction-level contributions were summed by order level; fractional contributions were calculated relative to total pathway abundance.

#### AI disclosure

Generative Large Language Models (LLMs) (ChatGPT, OpenAI, Dolphin AI) were used for language editing/revising and structural refinement of the manuscript. Generative LLMs were also consulted editorially to suggest further ideas for subsections of the Discussion section, as well as for statistical design ideas regarding the prevalence-aware analysis for Supplementary Figures S3 and S4. All experimental design, data analysis, statistical testing, figure generation, and scientific interpretation were conducted independently by the authors, and generative LLM editorial suggestions were vetted and further revised, if included.

## Results

### Taxonomic composition of the phylloplane epimicrobiome

Taxonomic profiling of metatranscriptomic reads using MetaPhlAn4 identified 83 microbial species spanning 27 taxonomic orders (predominantly bacterial, except for the two fungal orders Malasseziales and Eurotiales) in inoculated leaf samples (Figure 1). Across hostplant species, phylloplane (epimicrobiome) communities were dominated by members of Sphingomonadales and Propionibacteriales, which were also detected in the soil inoculum, although their relative abundance was reduced on *Nepenthes bicalcarata* (Figure 1A). Few bacterial orders exhibited host-specific distributions, including Bacillales (largely restricted to *Gossypium* spp.) and Acidobacteriales (restricted to *Nepenthes* spp.), though several taxa within microbial orders exhibited some host-specificity, e.g., Nitrosomonadales (*Methylophilus sp Leaf408* in *Gossypium,* and *Methylotenera oryzisoli* in *Beta* and *Nepenthes*). Certain microbial orders detected at low abundance in the inoculum (i.e., Lactobacillales and Bacteroidales) were not recovered from inoculated leaves (Figure 1A).

**Figure 1.**
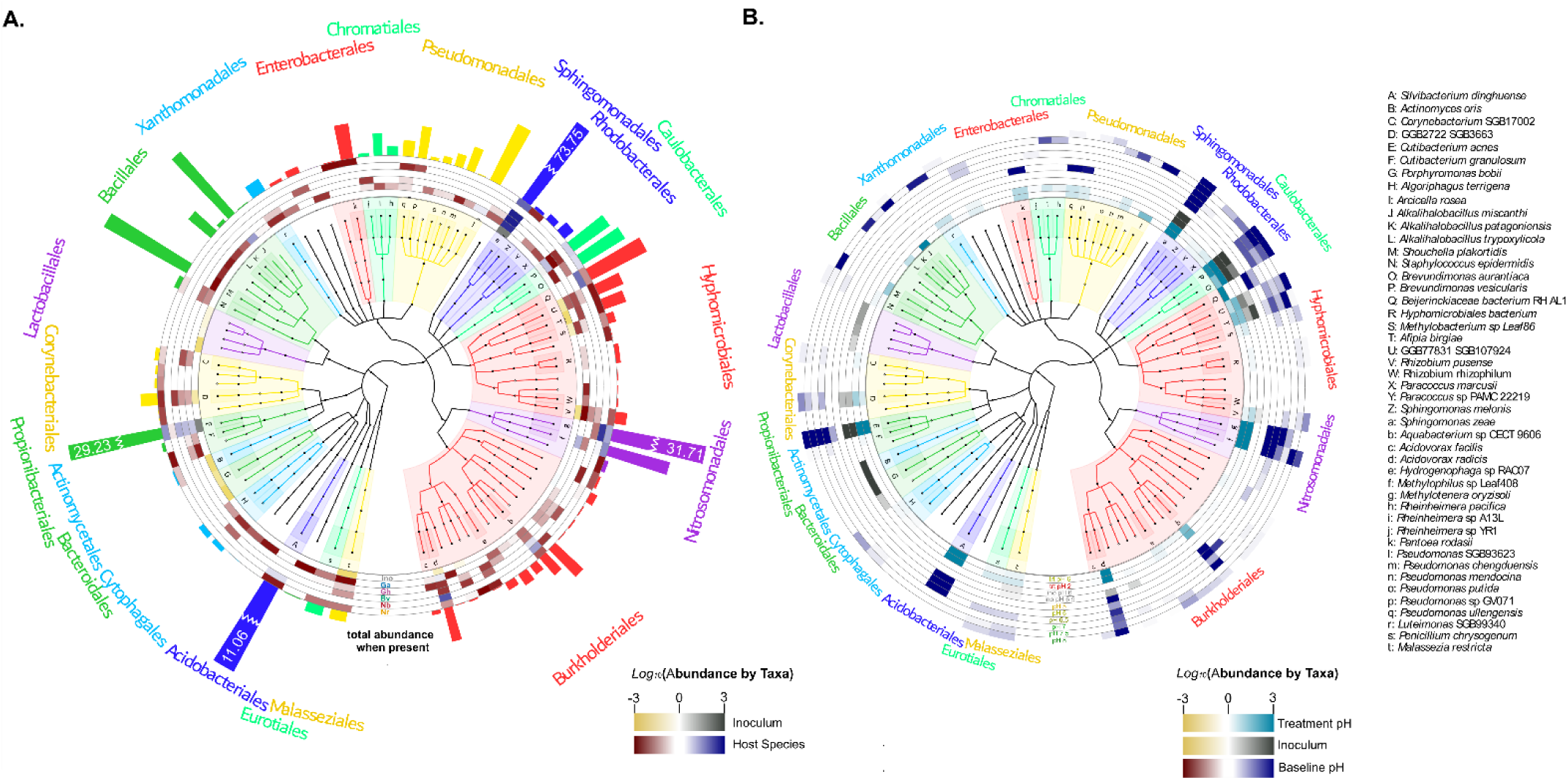
Phylogenetic distribution and relative abundance of phylloplane microbial taxa recovered from inoculated leaves. Taxonomic profiles were generated from metatranscriptome data using MetaPhlAn4. Microbial taxa with ≥0.01% relative abundance were visualized using GraPhlAn. The central circular dendrogram represents phylogenetic relationships among detected taxa based on the MetaPhlAn clade-specific marker gene reference tree. Microbial orders are labeled along the outer perimeter, and selected taxa are annotated to species level. Species-level classification was determined using MetaPhlAn4 clade-specific marker genes derived from genome-resolved phylogeny. Color intensity within heatmap rings represents log10-transformed relative abundance (see legend). **(A)** Hostplant-associated abundance patterns. The innermost ring indicates presence/absence of taxa in the soil inoculum (not introduced to leaves). Subsequent rings represent taxa detected on inoculated leaves of each hostplant species: Ga, *Gossypium arboreum*; Gh, *Gossypium hirsutum*; Bv, *Beta vulgaris*; Nb, *Nepenthes bicalcarata*; Nr, *Nepenthes rafflesiana*. The outermost bar plot shows total relative abundance of each taxon across all hostplant treatments. For four highly abundant orders (Sphingomonadales, Nitrosomonadales, Propionibacteriales, and Acidobacterales), bar lengths were truncated for visualization clarity; actual relative abundance values are indicated numerically. **(B)** pH-associated abundance patterns. Heatmap rings represent relative abundance of taxa across experimental pH treatments applied after treatment (pH 6.5 and pH 2), inoculum pH values measured at the end of the experiment, and baseline phylloplane pH values measured prior to pH manipulation. Rings corresponding to baseline pH are ordered from lowest to highest pH.

To examine associations with leaf surface chemistry, taxa were visualized across the two experimental pH treatments and along the gradient of baseline phylloplane pH measured from each leaf (Figure 1B). While experimental pH treatments applied after inoculation showed limited qualitative differences in taxon occurrence, baseline pH corresponded to shifts in the distribution of several orders; for example, Acidobacteriales were primarily detected on hostplants with moderately acidic baseline pH (5-6.5), whereas Caulobacterales were more frequently observed on neutral to slightly alkaline leaf surfaces (6.5-8; Figure 1B). Consistent with inoculum-to-leaf filtering, Shannon diversity differed between inoculum and leaf samples for the taxonomic profiles (Kruskal-Wallis χ² = 8.36, P = 0.0038), whereas Inverse Simpson diversity did not (P = 0.42), indicating turnover primarily among low-abundance taxa.

### Hostplant identity drives community structure, with host-dependent taxonomic responses to pH treatment

Constrained ordination of inoculated leaf samples revealed that hostplant species significantly structured microbial community composition at both taxonomic and functional levels (Figure 2). For taxonomic profiles (MetaPhlAn4), the RDA model explained 64.27% of total variance (constrained inertia = 0.316 of total inertia = 0.492), with RDA1 and RDA2 accounting for 45.96% and 13.98% of total variation, respectively (Figure 2A). Permutation tests indicated a strong hostplant effect (F₄,₁₈ = 5.67, P = 0.001). Although the experimental pH treatment applied after inoculation did not show a significant main effect (F₁,₁₈ = 0.62, P = 0.561), the hostplant species × treatment interaction was significant (F₄,₁₈ = 2.27, P = 0.029), indicating that taxonomic responses to the pH treatment differed among host species.

**Figure 2.**
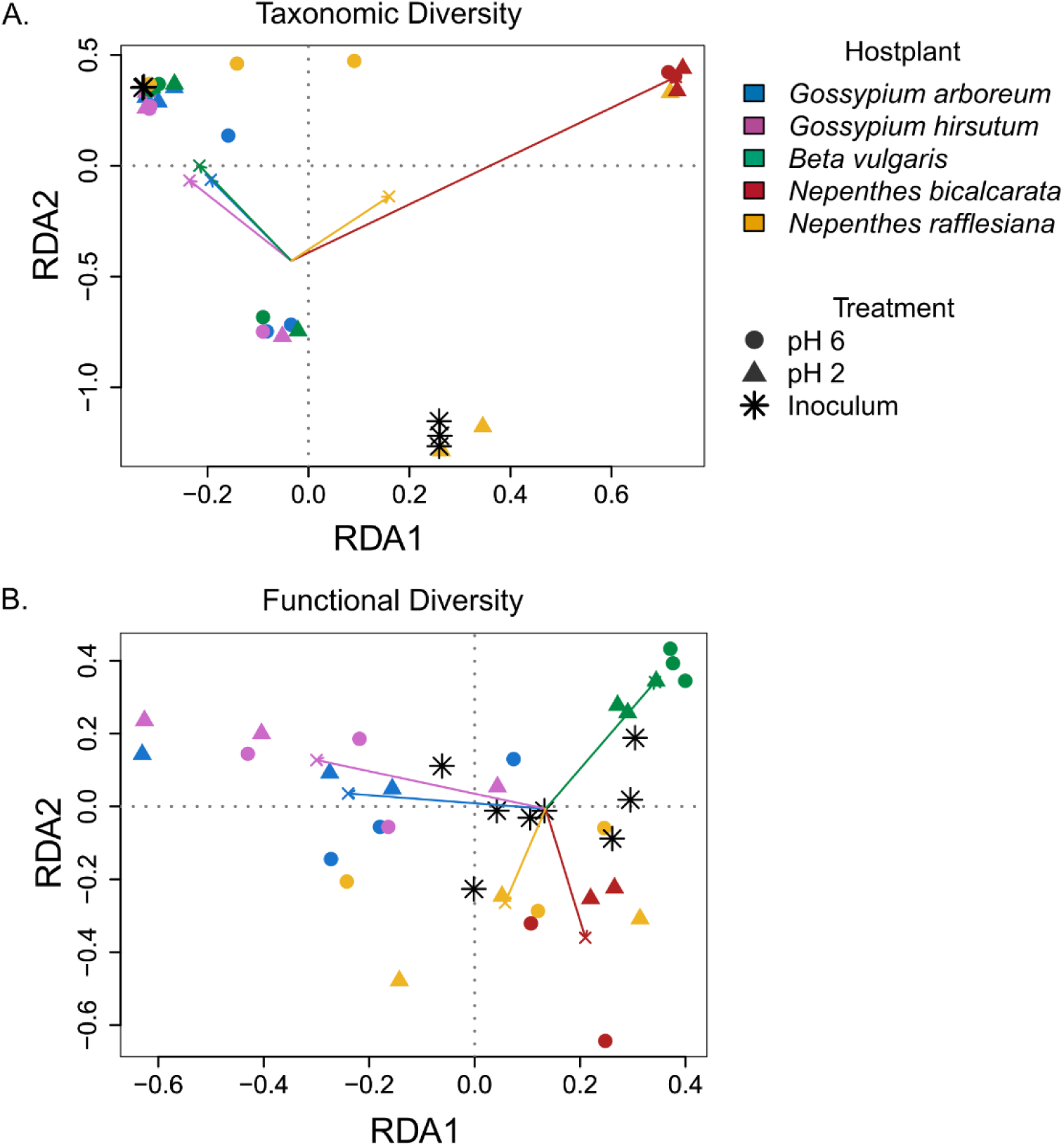
Hostplant identity structures phylloplane community composition at taxonomic and functional levels. **(A)** Redundancy analysis (RDA) of taxonomic composition based on MetaPhlAn4 species-level relative abundance profiles from inoculated leaf samples, constrained by hostplant species, experimental pH treatment (pH 6.5 vs pH 2), and their interaction. Points represent individual samples colored by hostplant species and shaped by treatment. The first two constrained axes (RDA1 and RDA2) explain 45.96% and 13.98% of total variance, respectively (71.52% and 21.75% of constrained variance). Hostplant species significantly explained community structure (permutation test, P = 0.001), and a significant host × treatment interaction indicated host-dependent responses to the pH treatment (P = 0.029), whereas the treatment main effect was not significant (P = 0.561). **(B)** RDA of functional composition based on reaction-level abundance profiles (HUMAnN3), constrained by the same model terms. The first two constrained axes explain 23.38% and 9.79% of total variance (44.27% and 18.54% of constrained variance). Hostplant species significantly structured functional profiles (P = 0.001), while treatment and the host × treatment interaction were not significant (P = 0.589 and P = 0.410, respectively).

Functional profiles based on reaction-level abundances (HUMAnN3) exhibited a similar dominant host effect but lacked evidence for host-dependent treatment responses (Figure 2B). The functional RDA explained 52.8% of total variance (constrained inertia = 0.069 of total inertia = 0.131), with RDA1 and RDA2 accounting for 23.38% and 9.79% of total variation, respectively. Hostplant species significantly structured reaction-level composition (F₄,₁₈ = 3.81, P = 0.001), whereas neither treatment nor the host × treatment interaction was significant (treatment: F₁,₁₈ = 0.80, P = 0.589; interaction: F₄,₁₈ = 1.03, P = 0.410). Together, these results indicate that host identity consistently structures both taxonomic and functional composition, whereas short-term experimental pH manipulation yields host-dependent taxonomic shifts that are not mirrored at the reaction-level functional resolution.

### Host identity dominates pH-associated patterns in functional composition across datasets

To test whether phylloplane pH independently structures microbial functional composition, we performed partial redundancy analyses (RDA) using reaction-level profiles (HUMAnN3) constrained by leaf surface pH, while conditioning on host identity (Figure 3). In inoculated leaves from the present experiment, baseline leaf pH (the value measured after inoculation, but before pH manipulation) explained only a small fraction of functional variation after accounting for host species (R² = 0.0349, adj. R² = 0.0108) and was not statistically significant (F = 1.36, P = 0.199). In contrast, baseline pH was significant in an unconditioned model that did not account for host identity (R² = 0.1055, adj. R² = 0.0711; F = 3.07, P = 0.005; Supplementary Figure S1A), indicating that pH-associated functional differences are largely embedded within broader host-associated variation.

**Figure 3.**
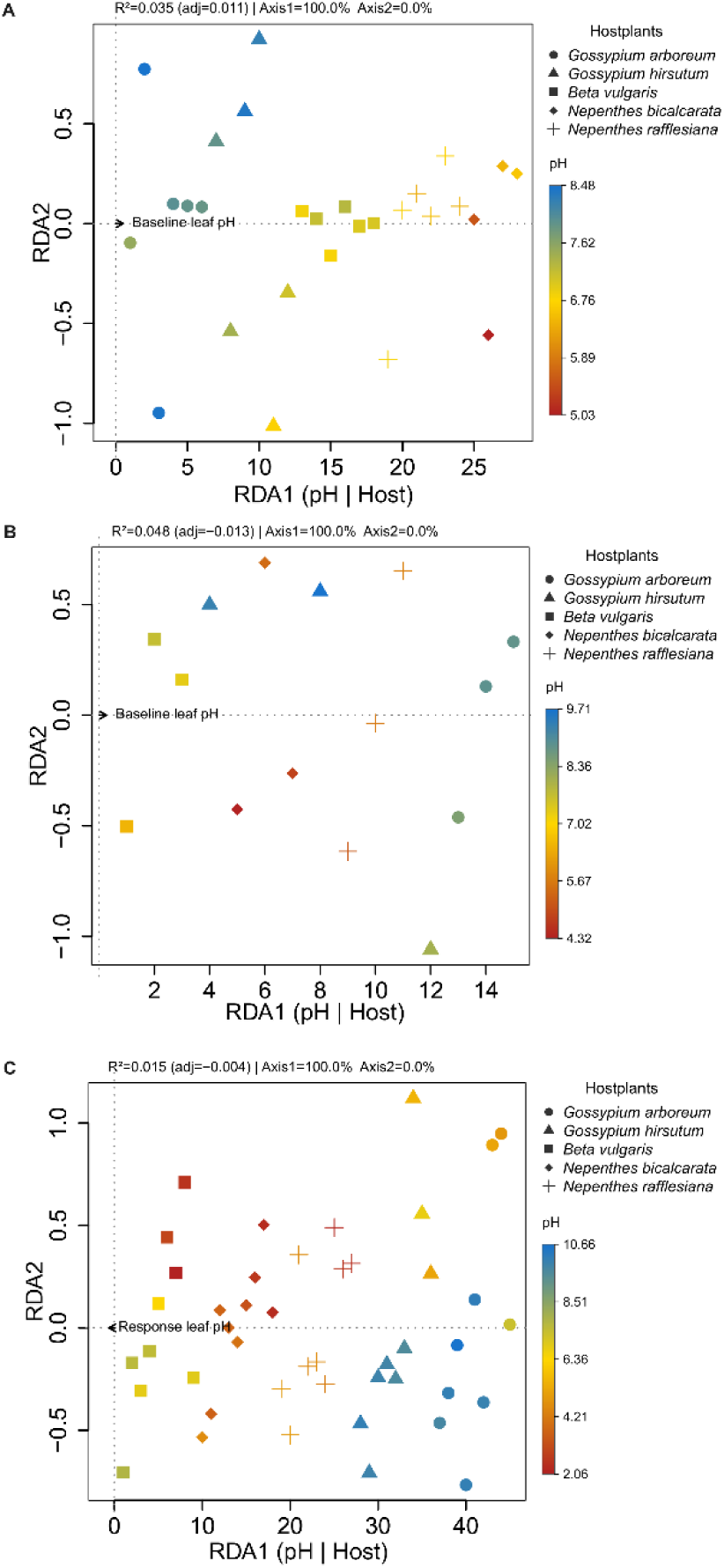
Leaf surface pH explains limited functional variation after accounting for host identity across experimental and external datasets. Partial redundancy analysis (RDA) of reaction-level functional profiles (HUMAnN3) constrained by leaf surface pH and conditioned on host species. Points represent individual samples colored by the pH gradient. All P-values were obtained using 999 permutations. **(A)** Inoculated leaves from the present study. Baseline leaf surface pH (measured after inoculation and prior to experimental pH manipulation) did not significantly explain functional composition after conditioning on host identity (F = 1.36, P = 0.199; R² = 0.0349, adj. R² = 0.0108). **(B)** Uninoculated leaves from a previous published dataset (NCBI GEO accession GSE281272). Baseline pH measured on dry leaf surfaces (control, no spray) was not associated with functional composition after conditioning on host species (F = 0.84, P = 0.672; R² = 0.0481, adj. R² < 0). **(C)** Uninoculated leaves (NCBI GEO accession GSE281272) subjected to pH spray treatments (pH 6.5, pH 4, pH 2). Response pH measured 5 min after spraying did not significantly explain functional composition after conditioning on host species (F = 0.81, P = 0.869; R² = 0.0152, adj. R² < 0).

We then evaluated whether pH-function relationships were detectable in the microbial portion retrieved from a previous independent, uninoculated whole-leaf transcriptome dataset for the same hostplant species (NCBI GEO accession GSE281272) [17]. In control samples collected from dry, unsprayed leaf surfaces, baseline phylloplane pH was not associated with functional composition after conditioning on host identity (F = 0.84, P = 0.672; Figure 3B). Similarly, following short-term pH spray treatments (pH 6.5, 4 or 2), response pH measured 5 min after spraying did not significantly explain functional composition after conditioning on host identity (F = 0.81, P = 0.869; Figure 3C). Consistent with host-structured sampling in this parallel experimental dataset, unconditioned models evaluated with permutations restricted within host species likewise did not detect significant pH associations (Supplementary Figure S1B, S1C). Together, these analyses indicate that host identity is the dominant axis of functional differentiation, and that leaf surface pH, whether baseline or spray-induced, explains little additional variation in reaction-level functional composition once accounting for host effects.

To quantify the magnitude of host-associated filtering relative to the source community, we calculated Bray-Curtis distances between each leaf sample and the inoculum based on reaction-level profiles (HUMAnN3). Distance to inoculum differed significantly among hostplants (Kruskal–Wallis χ² = 12.64, P = 0.013; ANOVA host effect: F₄,₁₈ = 4.29, P = 0.013; Supplementary Figure S2), whereas neither treatment nor the host × treatment interaction was significant (P > 0.62). Pairwise comparisons indicated that *Beta vulgaris* harbored communities significantly closer to the inoculum than *Gossypium arboreum* (P = 0.0028) and *G. hirsutum* (P = 0.0323). These results demonstrate that host species differ in the extent to which they filter or restructure the inoculum-derived functional pool.

### Host-associated filtering restructures the reaction repertoire relative to the inoculum

To quantify how leaf colonization reorganizes functional potential relative to the source inoculum, we analyzed HUMAnN3-derived reaction features (RXNs) as prevalence-filtered presence-absence matrices and visualized curated intersections across inoculated leaf epimicrobiomes, uninoculated whole-leaf datasets, and the soil inoculum (Figure 4A).

**Figure 4.**
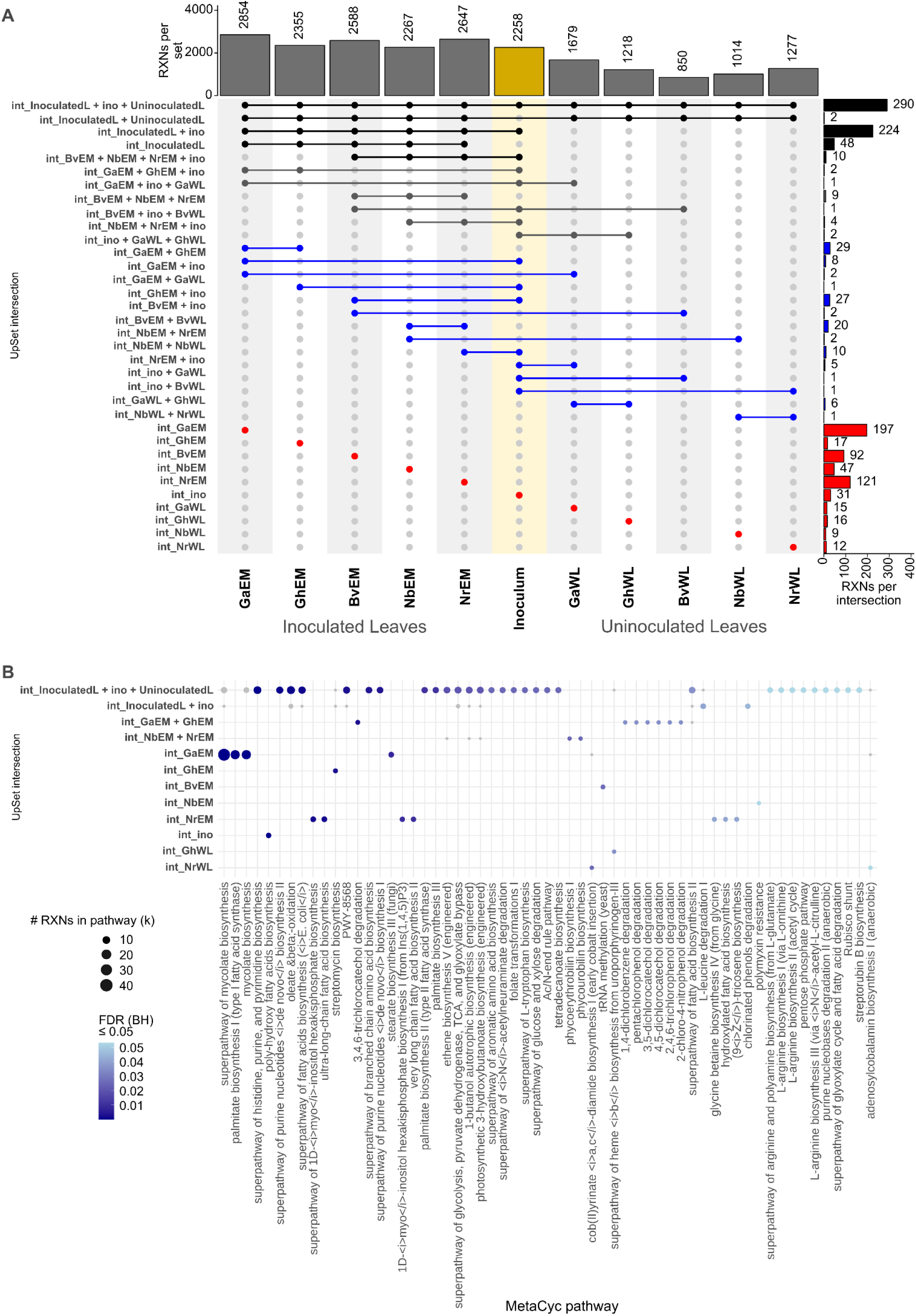
Prevalence-aware reaction intersections across inoculated leaf epimicrobiomes, uninoculated whole-leaf datasets, and the soil inoculum reveal shared and context-specific functional repertoires and enriched MetaCyc pathways. **(A)** UpSet plot summarizing overlapping intersections (int_) of HUMAnN3-derived reaction features (RXNs) across inoculated leaf epimicrobiome groups (GaEM, GhEM, BvEM, NbEM, NrEM), the soil inoculum (ino), and uninoculated whole-leaf datasets from an independent study of the same host species (GaWL, GhWL, BvWL, NbWL, NrWL). Reaction abundances were CPM-normalized using HUMAnN3 and converted to presence-absence using a prevalence-aware criterion (reaction considered present if CPM ≥ 1 in ≥66% of samples within a group; at least one replicate required). Top bars indicate the total number of reactions detected per group (set size), and right bars indicate the number of reactions within each displayed intersection (intersection size). Dots and connecting lines define the group combination corresponding to each intersection. Displayed intersections include group-specific reactions and selected shared combinations. **(B)** MetaCyc pathway enrichment analysis for reaction sets corresponding to the intersections shown in panel A (restricted to intersections containing ≥10 reactions). Reactions were mapped to MetaCyc pathways using reaction-to-pathway annotations, and enrichment was assessed using a one-sided hypergeometric test with the background defined as the union of all annotated reactions across tested intersections. P-values were adjusted using the Benjamini–Hochberg method. Points indicate enriched pathways within each intersection; point size reflects the number of reactions from that intersection mapping to the pathway (k), and point color indicates FDR-adjusted significance (q ≤ 0.05). Only pathways with (q ≤ 0.05) are shown.

Across all groups, a substantial shared reaction core was detected, indicating that leaf-associated communities draw from a largely conserved functional pool. The largest intersection comprised reactions detected across all inoculated hosts, uninoculated datasets, and the inoculum, consistent with a common metabolic backbone. In parallel, structured context-specific subsets were evident. Inoculated leaf epimicrobiomes retained many inoculum-derived reactions while exhibiting host-associated subsets not detected in the inoculum, indicating host-associated filtering beyond simple inoculum retention.

Under the primary analysis detection-based threshold (Figure 4A), overall reaction richness was broadly comparable among inoculated hosts. In contrast, uninoculated whole-leaf transcriptomes exhibited reduced reaction counts when stricter prevalence filtering was applied (Supplementary Figure S3). This reduction likely reflects lower microbial biomass and sequencing depth in plant transcriptomes not specifically designed to capture microbial signal, rather than true absence of functional capacity. Importantly, despite this quantitative reduction, the dominant shared intersections and overall structural relationships among groups remained qualitatively consistent under stricter filtering (Supplementary Figure S3A), supporting robustness of the core functional overlap.

Prevalence-aware estimates of group-specific reaction fractions (Supplementary Figure S4A), calculated as the proportion of reactions detected exclusively in a single host relative to its total repertoire, further showed that no host was dominated by a large proportion of unique reactions. Differentiation therefore primarily reflects restructuring of a shared functional pool rather than acquisition of novel metabolic capabilities.

Consistent with this interpretation, reaction repertoire turnover relative to the inoculum (Jaccard distance; Supplementary Figure S4B) differed among host species. The epimicrobiome of inoculated leaves exhibited moderate restructuring of the inoculum-derived repertoire, whereas uninoculated whole leaf epimicrobiomes showed higher apparent turnover under stricter filtering (SupplementaryFigure S4C, S4D), consistent with sparse microbial detection rather than ecological divergence alone.

Together, these analyses demonstrate that host identity reshapes the reaction repertoire through selective retention and filtering of a largely shared functional backbone, and that this conclusion is robust to filtering stringency.

### Enriched pathway classes reflect structured functional filtering across intersections

To determine whether intersection-specific reaction sets were functionally structured, we mapped reactions to MetaCyc pathways and performed enrichment analyses for intersections containing ≥10 reactions (Figure 4B).

Enriched pathways spanned amino acid biosynthesis, nucleotide metabolism, central carbon metabolism, lipid biosynthesis, and aromatic compound degradation. The largest shared intersections were enriched for broadly conserved biosynthetic and housekeeping pathways, consistent with a common functional backbone across leaf-associated microbiomes. Smaller host-specific intersections were enriched for more specialized metabolic processes of stress-associated, envelope-related, and carbon-conservation pathways, indicating structured filtering of functional potential (retention of particular trait combinations) rather than stochastic loss. Importantly, enrichment patterns were not driven by single taxa but reflected composite contributions across multiple bacterial orders.

Applying stricter prevalence thresholds (Supplementary Figure S3B) reduced the total number of reactions in some intersections-particularly in uninoculated transcriptomes but preserved the qualitative pattern of shared-core enrichment and host-associated pathway structuring. These results indicate that pathway-level differentiation is robust to filtering stringency and not driven by low-prevalence reactions.

### Host-associated pathway differences are detectable relative to the inoculum baseline, whereas treatment effects within leaves are limited

To test for quantitative shifts in pathway abundance, we modeled MetaCyc pathway profiles inferred from HUMAnN3 reaction abundances using MaAsLin3 (Figure 5). Relative to the inoculum baseline, multiple pathways differed significantly across host-treatment groups. These included biosynthetic pathways (e.g., amino acid and nucleotide biosynthesis), cell envelope-associated processes (e.g., peptidoglycan maturation), and central carbon metabolism pathways (e.g., pentose phosphate pathway and glyoxylate cycle). Several aerobic respiration-associated pathways exhibited reduced relative abundance on leaves compared to inoculum (Supplementary dataset S1), whereas other biosynthetic and carbon metabolism pathways showed positive shifts, consistent with altered metabolic states following leaf colonization.

**Figure 5.**
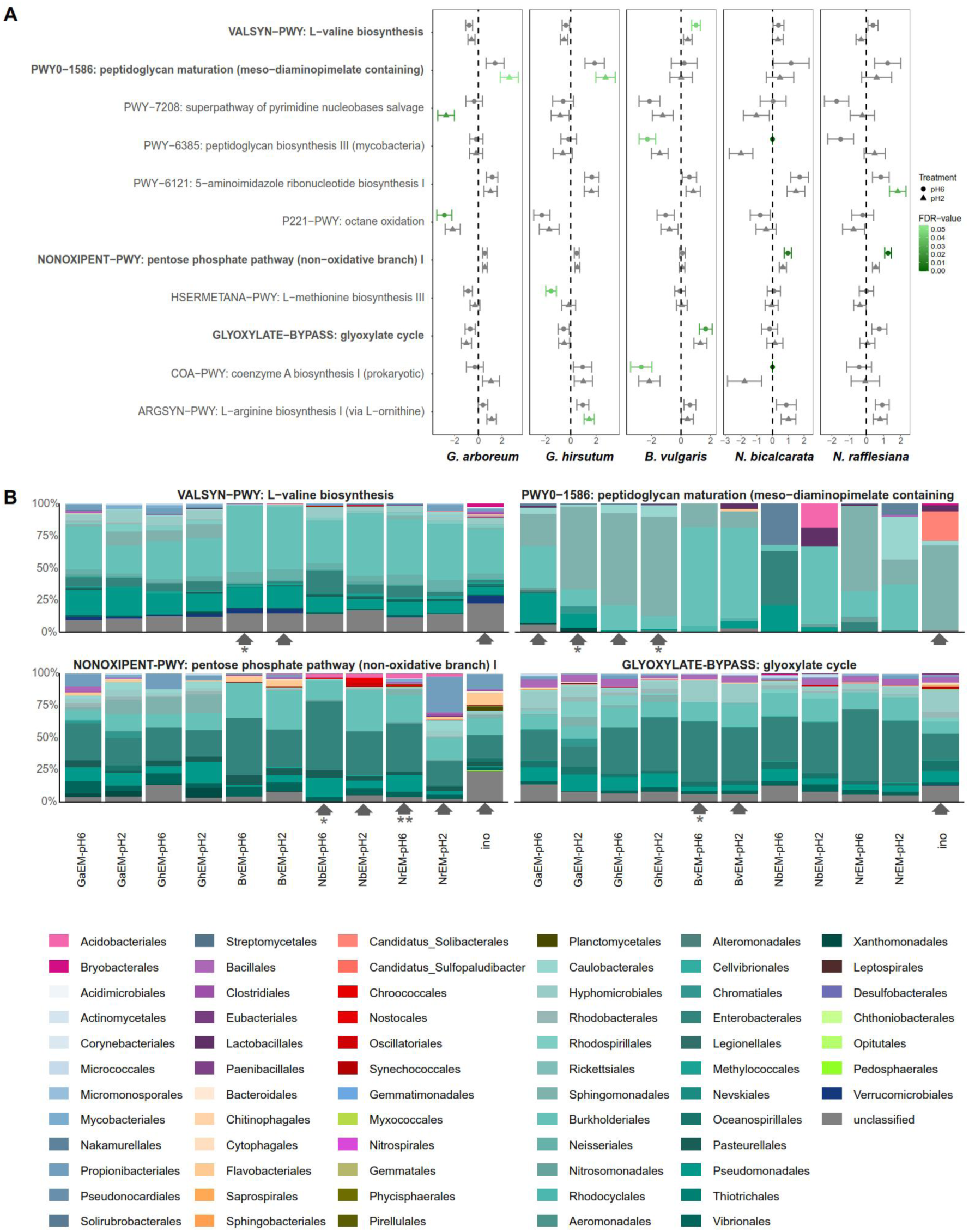
Host-associated shifts in microbial functional pathways and their taxonomic contributors. **(A)** Differentially abundant MetaCyc pathways across hostplant-treatment groups relative to the inoculum baseline. Pathway abundances were taken from HUMAnN3 MetaCyc pathway profiles and tested with MaAsLin3 using hostplant-treatment group as the predictor, with inoculum as the reference level. For each hostplant epimicrobiome (EM) (*Gossypium arboreum*, *G. hirsutum*, *Beta vulgaris*, *Nepenthes bicalcarata*, and *N. rafflesiana*), points show the estimated effect size (β coefficient ± SE) for each hostplant-pH group (pH 6.5, pH 2) relative to inoculum; positive coefficients indicate higher pathway abundance on leaves than inoculum and negative coefficients indicate lower abundance, on the *log*-transformed scale used by MaAsLin3. Only pathways significant in at least one host-treatment comparison after FDR correction (q ≤ 0.05) are shown. Names of selected pathways of interest are in bold. **(B)** Microbial taxonomic order contributions to representative significant pathways (examples shown: VALSYN-PWY, PWY0-1586, NONOXIPENT-PWY, and GLYOXYLATE-BYPASS). These pathways were selected because they show a significant pattern of expression in at least one of the hostplant taxa, highlighting how different hostplants are selecting for different microbial functions. Stacked bars represent the proportional contribution of each microbial order to total pathway abundance for each host-treatment combination and the inoculum (order-stratified HUMAnN3 output collapsed to order level and converted to percentages per pathway). “Unclassified” denotes reads not assigned to an order. Gray arrows highlight the turnover of microbial contribution to that feature between the two pH treatments (6.5, 2), within a given hostplant species, that shows significant differential abundance compared to inoculum. Asterisks beneath the arrows indicate host-treatment groups significantly different from inoculum after FDR correction (q ≤ 0.05).

In contrast, when analyses were restricted to inoculated leaf samples, no pathways were detected as significant for the host × treatment interaction. Multivariate analyses similarly provided no evidence that hostplant:treatment structured pathway composition (PERMANOVA by margin: R² = 0.095, F = 0.707, P = 0.852), and dispersion did not differ among host, treatment, or host×treatment groups (all betadisper P > 0.21).

Thus, consistent with reaction-level intersection analyses (Figure 4), host-associated filtering relative to inoculum is detectable at the pathway-abundance level, whereas short-term pH manipulation does not induce consistent functional reorganization across hosts.

### Taxonomic contributors to key pathways differ across host contexts

To link pathway-level shifts to community composition, we quantified order-level contributions to representative pathways that differed relative to inoculum (Figure 5B). Contributions to the non-oxidative pentose phosphate pathway (NONOXIPENT-PWY) and glyoxylate cycle (GLYOXYLATE-BYPASS), for example, were distributed across multiple bacterial orders, with relative contributions varying among hosts and between leaves and inoculum.

Across pathways, expressed functional profile on leaves reflected composite contributions from multiple microbial orders rather than dominance by a single lineage (Supplementary Figure S5). These results indicate that hostplant filtering alters not only which taxa persist (Figures 1-2), but also the taxonomic sources contributing to pathway-level expression pattern.

### Host-associated GO-term differences mirror pathway-level patterns

To evaluate shifts in expressed functional annotations, we tested GO-term abundances derived from HUMAnN3 and visualized significant terms alongside their microbial contributors (Figure 6). Multiple GO terms differed between host-treatment groups and the inoculum baseline, spanning functions related to translation and protein synthesis, redox and energy metabolism, transport, and motility-associated processes. Many significant terms exhibited reduced abundance on leaves relative to inoculum (Supplementary Dataset S1), consistent with colonization-associated restructuring of functional activity.

**Figure 6.**
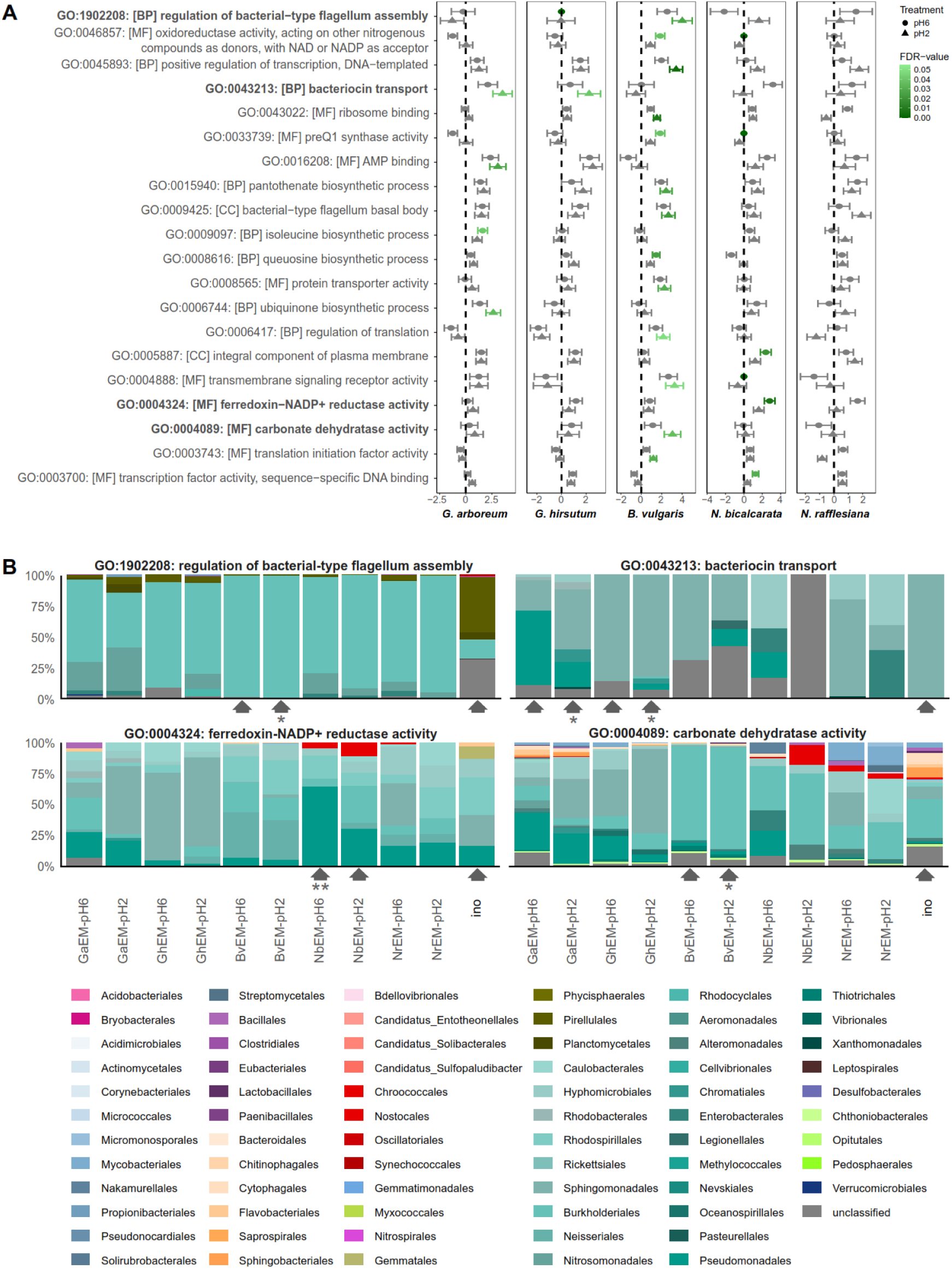
Host-associated differences in GO-annotated functions and their taxonomic contributors. **(A)** Differentially abundant Gene Ontology (GO) terms across hostplant-treatment groups relative to the inoculum baseline. GO-term abundances were generated from HUMAnN3 output by regrouping gene-family contributions to GO terms and renormalizing to counts per million (CPM), then tested with MaAsLin3 using hostplant-treatment group as the predictor and inoculum as the reference level. For each hostplant epimicrobiome (EM) (*Gossypium arboreum*, *G. hirsutum*, *Beta vulgaris*, *Nepenthes bicalcarata*, and *N. rafflesiana*), points show the MaAsLin3 estimated effect size (β coefficient ± SE) for pH 6.5 and pH 2 groups relative to inoculum; positive coefficients indicate higher GO-term abundance on leaves than in the inoculum and negative coefficients indicate lower abundance, on the log-transformed scale used by MaAsLin3. Point color denotes FDR-adjusted q-value. Only GO terms significant in at least one hostplant-treatment comparison after FDR correction (q ≤ 0.05) are shown. Names of selected GO terms of interest are in bold. **(B)** Microbial taxonomic order contributions to representative significant GO terms (examples shown: GO:1902208 regulation of bacterial-type flagellum assembly; GO:0043213 bacteriocin transport; GO:0004324 ferredoxin-NADP+ reductase activity; GO:0004089 carbonate dehydratase activity). These GO terms were selected because they show a significant pattern of expression in at least one of the hostplant taxa, highlighting how different hostplants are selecting for different microbial functions. Stacked bars represent the proportional contribution of each microbial order to total GO-term abundance for each hostplant-treatment group and the inoculum, based on the order-stratified HUMAnN3 output collapsed to order level and converted to percentages within each GO term. “Unclassified” denotes reads not assigned to an order. Gray arrows highlight the turnover of microbial contribution to that feature between the two pH treatments (6.5, 2), within a given hostplant species, that shows significant differential abundance compared to inoculum. Asterisks beneath the arrows indicate host-treatment groups significantly different from inoculum after FDR correction (q ≤ 0.05).

However, analyses restricted to inoculated leaf samples provided no evidence that hostplant:treatment structured GO-term composition (PERMANOVA by margin: R² = 0.109, F = 1.112, P = 0.307), and dispersion did not differ among groupings (all betadisper P ≥ 0.15). Order-level contribution analyses (Figure 6B) indicated that shifts in GO-term abundances reflect contributions from multiple bacterial orders, rather than dominance by a single lineage. Thus, host filtering reorganizes the taxonomic architecture of functional capacity without generating strong, uniform treatment-dependent shifts in expressed function.

## Discussion

Plant-associated microbiomes are shaped by both environmental conditions and host traits, yet the relative contribution of these forces in the phyllosphere has been an outstanding question. While soil and rhizosphere are well established as host-structured systems [1,2], aboveground epiphytic microbial communities are often viewed as primarily influenced by external abiotic factors such as temperature, moisture, and atmospheric deposition [3,4]. Here, we show that hostplant identity is the dominant axis structuring phylloplane microbiomes at both taxonomic and functional levels. Leaf colonization does not generate novel functional repertoires; rather, hosts reorganize a largely shared inoculum-derived reaction pool through selective retention and filtering. This restructuring is consistent across different analyses—from reaction-level intersections and pathway enrichment, to quantitative shifts in pathway and GO-term abundances—and is robust to prevalence-filtering thresholds. In contrast, short-term manipulation of leaf surface pH does not induce consistent community-wide functional reorganization once host effects are accounted for, suggesting that transient chemical perturbation is embedded within broader host-associated variation. Collectively, these findings position leaves not as passive substrates exposed to environmental fluctuation, but as active ecological filters that structure the taxonomic architecture and functional capacity of associated microbiomes. We additionally provide insight into how phylloplane bacteria may physiologically respond to different leaf environments and pH conditions at an organismal level.

### Host identity structures microbial composition and gene expression

Across taxonomic (MetaPhlAn4) and functional (HUMAnN3 reaction-level) profiles, hostplant species identity consistently explained the largest fraction of variation (Figures 1-2). This pattern was maintained across epimicrobiomes from experimentally inoculated leaves and microbiomes from the uninoculated whole-leaf dataset (Figures 3-4), indicating that host identity structures microbial gene expression across leaf-associated habitats. As a caveat to all functional profile results, metatranscriptomics quantifies transcript abundance rather than metabolic flux [26], thus our conclusions pertain to expressed functional potential rather than realized enzymatic activity rates. However, examining functional potential still provides coherent insight into the active microbial community, interpretable as a trait-based view of community assembly complementary to the taxonomic view.

Alpha-diversity patterns further support selective restructuring of microbial communities during leaf colonization. Shannon diversity differed significantly between inoculum and leaf-associated communities, whereas Inverse Simpson diversity did not, indicating that turnover occurred primarily among low-abundance taxa while dominant taxa remained comparatively stable. Because Shannon diversity is more sensitive to rare members of the community, this pattern suggests that community turnover was driven by changes in low-abundance taxa, with dominant taxa remaining more stable. Such asymmetric turnover is consistent with ecological filtering that constrains establishment of marginal taxa without disrupting core community structure [47].

We selected diverse hostplants representing a wide range of baseline phylloplane pH levels. Although we found a relationship between epimicrobiome composition and baseline pH, its explanatory power diminished when host identity was included in multivariate models (Figure 3; Supplementary Figure S1). This may indicate that pH does not operate as an independent abiotic driver, but rather as one component of a broader host-defined ecological regime. Future work with more closely related plant species that vary in phylloplane pH (or one hostplant species with strong intraspecific variation in the trait) may help better disentangle the effect of host identity from the effect of baseline pH per se, however, our current work represents a pivotal first step. Another consideration might be whether longer-term pH perturbations may have greater effects on the structure of the microbiome; previous work characterizing phylloplane pH buffering shows that after exposure to neutral water, the peak pH is reached within five minutes and then plateaus over the course of an hour [16]. While host-dominated structuring is well-established belowground [1,2], evidence for comparably strong host effects in the phyllosphere has been more context-dependent [5,20]. Our results extend this host-centric view to leaf-associated microbial gene expression landscapes under controlled colonization conditions. Further, our investigation is a prime example of the phytobiome concept: to fully understand a plant, biotic and abiotic features of its ecosystem and its interacting symbionts must be considered together in a holistic fashion [48]. In our case, plant-microbial interactions occur in the context of pH, an abiotic feature of the external environment, which is shaped by the plant [49], shapes the plant [17], shapes the microbes, and variously influences how plant and microbe interact with one another.

### Leaves restructure a shared environmental functional backbone

Reaction-level intersection analyses revealed a substantial functional core shared across inoculated leaf epimicrobiomes, the soil inoculum, and microbiomes from the uninoculated whole-leaf dataset (Figure 4A). The largest intersections comprised reactions detected across all hosts and contexts, indicating that leaf-associated microbiomes draw from a broadly conserved metabolic reservoir. Considering the arbitrary nature of the source inoculum, and the lack of microbial standardization for the uninoculated leaves, this metabolic reservoir may be environmentally ubiquitous. Host-associated differentiation therefore did not arise from the emergence of host-specific metabolic novelty. Instead, it reflected selective retention and differential representation of reactions within a shared environmental pool. Prevalence-aware estimates confirmed that unique reaction fractions were small across hosts, reinforcing that functional differentiation primarily involved restructuring of a common backbone (Supplementary Figure S4A, S4C). Importantly, stricter prevalence filtering reduced reaction counts, particularly in uninoculated transcriptomes with lower microbial biomass, but preserved the qualitative intersection topology (Supplementary Figure S3), indicating that the shared core structure is robust to filtering thresholds. This pattern aligns with broader evidence that microbial ecosystems exhibit high functional redundancy, whereby taxonomically distinct communities retain overlapping metabolic capabilities [21,22]. In the phyllosphere, host identity appears to reorganize this shared functional space rather than constructing discrete metabolic repertoires. As expected, inoculated leaves had many commonalities with the inoculum, but there was negligible convergence of unique functional features between inoculated and uninoculated leaves of the same host species (Figure 4A). We also identified different molecular pathways represented by a few intersections of samples, but mostly involving the epimicrobiomes (Figure 4B).

### Functional filtering is metabolically structured

Mapping reaction intersections to MetaCyc pathways revealed that host-associated reaction subsets were metabolically coherent rather than random assemblages (Figure 4B). Reactions shared across hosts and contexts were enriched for central biosynthetic and core metabolic pathways, including amino acid and nucleotide biosynthesis and central carbon metabolism, forming a conserved functional backbone associated with microbial persistence on leaves [3,21,22]. In contrast, smaller host-associated intersections were enriched for more specialized functions, including pathways linked to oxidative stress management, carbon conservation (e.g., glyoxylate bypass), and envelope-associated processes, suggesting structured shifts in expressed functional composition rather than random loss of which pathways are present.

These pathway-level shifts align with ecological expectations for microbes transitioning from a homogenous nutrient-rich slurry to heterogeneous, nutrient-limited, and oxidatively stressful phyllosphere microhabitats [3–5]. For example, enrichment of pentose phosphate pathway components is consistent with increased NADPH generation for redox balance under UV- and ROS-exposed leaf environments [3,7], while representation of glyoxylate-cycle reactions supports carbon-efficient metabolism under carbohydrate limitation typical of the phyllosphere [3]. Concomitantly, aerobic respiration-related pathways had relatively reduced abundance in phylloplane epimicrobiomes, again supporting the relative nutrient limitation on leaf surfaces in contrast to the soil-derived inoculum [50], possibly accompanied by shifts to alternative carbon substrates [51] and/or shifting from rapid growth to maintenance metabolism throughout the community [52]. Finally, the apparent failure of Lactobacillales to successfully colonize the phylloplane epimicrobiome further supports the nutrient limitation hypothesis, as this order is generally restricted to extremely nutrient-replete habitats [53]. Relevant to the likely heterogenous distribution of nutrients on the phylloplane, differential representation of cell-envelope and motility-associated functions suggests restructuring of traits linked to surface persistence, attachment, and microscale habitat navigation on leaves [3–5].

Taxonomic contribution profiles further suggest that these pathway-level differences are not driven by wholesale replacement of dominant lineages, but by redistribution of functional contributions across multiple bacterial orders (Figures 5B, 6B; Supplementary Figure S5). In several cases, lower-abundance orders contributed proportionally more to differentially abundant pathways on leaves relative to inoculum, consistent with turnover within functional guilds rather than loss of metabolic capacity. Such patterns align with theoretical and empirical evidence for widespread functional redundancy in microbial systems, in which environmental shifts reorganize taxonomic contributors while maintaining core metabolic capabilities [21,22]. Host-associated filtering therefore appears to alter who performs a function as opposed to altering whether the function exists.

Across hosts, however, the magnitude of functional restructuring was not uniform. *Gossypium* leaf communities diverged most strongly from the inoculum at the reaction level (Figures 3-4; Supplementary Figure S2), indicating comparatively stronger host-associated reorganization of the functional repertoire. Although phylloplane pH did not independently explain this divergence in multivariate models, *Gossypium* is characterized by rapid alkalinization, mineral-rich secretions, and strong buffering capacity relative to *Beta* and *Nepenthes* [14,16,17,54,55]. It is also notable that the one hostplant species whose baseline pH levels were consistently neutral and closest to the original pH of the inoculum (mean pH 7.03 and pH 6.87, respectively) is also the species whose epimicrobiome is the least diverged from the inoculum: *Beta vulgaris*. While these results are suggestive of the possibility that the degree of pH modification determines the degree of ecological filtering by hostplants, it is also probable that a combination of phylloplane traits (e.g., ionic composition, cuticle chemistry, surface microstructure) act in concert with phylloplane pH to characterize the differences between hosts. In any case, strong ecological filters like *Gossypium* do not generate novel metabolic capacity, but rather more strongly restrict which subsets of a shared environmental pool persist and are expressed, yielding greater functional divergence relative to other hosts.

### Leaf-associated assembly effects outweigh short-term abiotic perturbation

Relative to the inoculum baseline, colonization of leaf surfaces produced detectable shifts in pathway representation (Figure 5), including changes in central carbon and biosynthetic metabolism. These results indicate transcriptional restructuring following leaf colonization. In contrast, short-term experimental pH manipulation did not induce consistent host × treatment effects among inoculated leaves. Neither pathway-level nor GO-term analyses revealed robust treatment-driven reorganization, and multivariate tests similarly lacked evidence of treatment-associated structuring at the community level (Figures 5-6; Supplementary Dataset S1).

Although pH is often considered a master variable in microbial ecology [8], transient abiotic perturbations may be insufficient to override host-imposed functional organization once communities are established. This resistance to short-term disturbance may reflect functional redundancy within the shared metabolic backbone, enabling communities to buffer transient environmental fluctuations [47]. It is also possible that the resistance of the functional microbial community to the pH perturbation may be due to the ability of hostplants to buffer against external change. However, we expected stronger functional preservation across treatments for *Gossypium*, which rapidly neutralizes pH 2 sprays (within 5 minutes), and a lack of functional preservation for *Beta*, which completely lacks this buffering ability [17]. So, the lack of substantial treatment-driven reorganization in *Beta* as well speaks to the precedence of inherent microbial metabolic causes as opposed to an effect of hostplant mediation in this case.

### Microbes respond to phylloplane pH at an organismal level

While we are unable to disentangle the effects of phylloplane pH from the effects of host identity in terms of community-level functional responses, we do have several points of evidence implicating the role of pH (both host-mediated and externally imposed) on the microbiome at finer scales. While taxonomic filtering may be due to a combination of host-specific traits across these disparate hostplants, it is notable that genes corresponding to the genera *Alkalihalobacillus* and *Shouchella* (Bacillales) only appear on the characteristically alkaline *Gossypium* phylloplane in our epimicrobiome experiment (Figure 1A). These two bacterial genera contain obligate alkaliphiles described from extreme soda lake environments [56], indicating that *Gossypium* leaves create extreme conditions akin to these larger-scale habitats. Further, these Bacillales taxa are the only detectable microbes that we observe as present in the pH 6.5 treatment whilst being absent at pH 2 (Figure 1B). Thus, while the short-term pH perturbation of the acid spray may not have sweeping effects on community assembly, it can impact the expression of particularly pH-sensitive community members, such as these low-abundance extremophiles. The mildly acidic or neutral environments of *Beta*, *Nepenthes*, and even the inoculum slurry are either unsuitable to the survival of these extremophiles, or more likely, suppress their activity. The acidic spray led to an artificially neutral *Gossypium* environment; apparently enough to diminish its baseline suitability for alkaliphiles. On the other end of the spectrum, despite being one of the four most abundant taxa, one taxon in Acidobacteriales putatively identified as the obligately acidophilic *Silvibacterium dinghuense* [57] was restricted to *Nepenthes* epimicrobiomes, only being expressed on baseline phylloplane pH levels below 6.5 (Figure 1A,B).

Fine-scale pH effects are further apparent at the organismal level, as observed in our pathway and GO term analyses. While we did not find treatment-driven functional reorganization at the community level, we did observe compelling trends suggesting responses both to external pH and host pH regulation. Particularly notable is the differential abundance of the peptidoglycan maturation pathway; in *Gossypium*, it is relatively more abundant at pH 2 whereas the opposite trend (though not significant) is observed in *Nepenthes,* and entirely low abundance for *Beta* (Figure 5). As peptidoglycan remodeling is one possible response to alkaline or acidic stress that bacteria use to maintain intracellular ionic homeostasis [58], it is biologically sensible that expression of this pathway may be more active on phylloplanes with extreme baseline pH levels than on the neutral phylloplane of *Beta*. Further, if the peptidoglycan maturation pathway is activated in response to the short-term pH perturbation, it is reasonable that the direction of its shift in expression would be opposite on alkalinizing vs. acidifying host phylloplanes. Notably, a GO term related to carbonate dehydratase activity was observed to be specifically associated with the *Beta* phylloplane environment, and more so at the pH 2 treatment (Figure 6). Carbonate dehydratases function in buffering extracellular environments [59], thus microbes on *Beta vulgaris* leaves, which is a host that lacks the ability to buffer against pH 2 sprays [17], may rely on these genes to shield themselves from the pH perturbation. The other hostplant species either rapidly neutralize the external pH spray *(Gossypium)* or cultivate more acid-tolerant microbes prior to the pH pulse (*Nepenthes*), diminishing the need for the microbes to self-buffer.

Finally, our data reveals surprising bacterial responses that merit future investigation. Regarding taxonomic filtering, results may point to pH preferences of important leaf-associated taxa, such as potentially beneficial methylotrophs in Nitrosomonadales [60–62]. In our study, different individual Nitrosomonadales taxa occurred on different hostplants differing in baseline pH, i.e. putative *Methylophilus* sp. Leaf408 on characteristically alkaline *Gossypium* and putative *Methylotenera oryzisoli* on neutral to acidic *Beta* and *Nepenthes* (Figure 1). All species-level taxonomic classifications in our study are tentative, but this potential niche differentiation is noteworthy, nonetheless. Returning to the functional implications of our pathway and GO term analysis, the trends of the bacteriocin transport GO term match the general pattern of the peptidoglycan pathway (Figure 6). Bacteriocin plays a role in bacteria-bacteria competition [63,64]. A controlled experiment on nectar microbes revealed that more extreme pH conditions (in their case, acidic) enhanced competitive interactions within microbiomes relative to neutral pH [65]. We found more responsive bacteriocin transport on the *Gossypium* phylloplane, suggestive of more extreme baseline pH (in our case, alkaline) enhancing competitive interactions. The neutral *Beta* phylloplane showed no apparent bacteriocin transport activity in its epimicrobiome. Instead, microbes on *Beta vulgaris* appeared to express genes related to flagella, which may indicate more active movement on this relatively permissive environment. This could also indicate that *Beta* is less effective at suppressing pathogenicity, as Type III secretion systems are derived from flagella [66,67], and we also see heightened activity of GO terms like “protein transporter activity” and “transmembrane signaling receptor activity” on *Beta*, especially when exposed to pH 2 (Figure 6). Future work should probe bacterial physiology in different host pH environments, and link microbial activity to hostplant fitness.

## Conclusions

In summary, hostplant identity is the dominant axis structuring both taxonomic composition and functional gene expression of microbial communities on the phylloplane; pH, in contrast, causes finer-scale physiological responses at the organismal level. Leaf-associated communities do not assemble entirely novel metabolic repertoires; rather, hosts selectively reorganize a shared environmental functional backbone. This restructuring is metabolically coherent and taxonomically distributed, consistent with functional redundancy and trait-based assembly. *Gossypium* imposes comparatively stronger reorganization, possibly reflecting a combination of phylloplane traits acting together along with pH. In contrast, short-term pH perturbation does not override host-structured functional organization at the community level, suggesting resilience of the leaf-associated microbial metabolic landscape. Together, these findings position leaves as active ecological filters that influence the distribution and expression of microbial functional capacity within the phytobiome. This work has also unveiled fine-scale responses of certain microbial taxa to hostplant traits and external pH, which calls for future controlled experiments directly probing the physiology of individual isolates, using techniques such as metabolomics, stable isotope labelling, and gene knockouts.

## Acknowledgements

We acknowledge Scott DiLoreto and Adam Bettinger for help with rearing plants in the greenhouse and growth chamber, as well as help from John Fulginiti and Elise Elizondo in preliminary methods testing. We also thank Mario Laterrière, Nicholas Panchy, Chloe Drummond, and Adam Rork for bioinformatics advice, and Kevin Hockett, Kylie Bocklund, William King, Lauren Sullivan, Cara Haney, Talia Karasov, Tess Grainger, members of the Plant Resilience Institute, and the Gilbert Lab for helpful discussions. We also thank Johan Leveau for a helpful comment on a much earlier version of this manuscript, which helped us significantly expand our scope. This work was supported by the Agriculture and Food Research Initiative - Education and Workforce Development program, Project #2019-67012-29872 and #2019-67012-37587, from the U.S. Department of Agriculture’s National Institute of Food and Agriculture. This work was supported by the USDA National Institute of Food and Agriculture and Hatch Appropriations under Project #PEN04974 and Accession #7006543.

## Author contributions

KJG and TR initially conceived of the idea, KJG performed greenhouse/growth chamber experiments and conducted RNA extractions for both the phylloplane and whole-leaf experiments, JBF and CLG performed bioinformatics for both experiments as well as all statistical analyses. JBF and KJG wrote the initial draft, and JBF, CLG, TR, and KJG contributed to writing the final version of the manuscript.

## Supplementary Figures

**Supplementary Figure S1.**
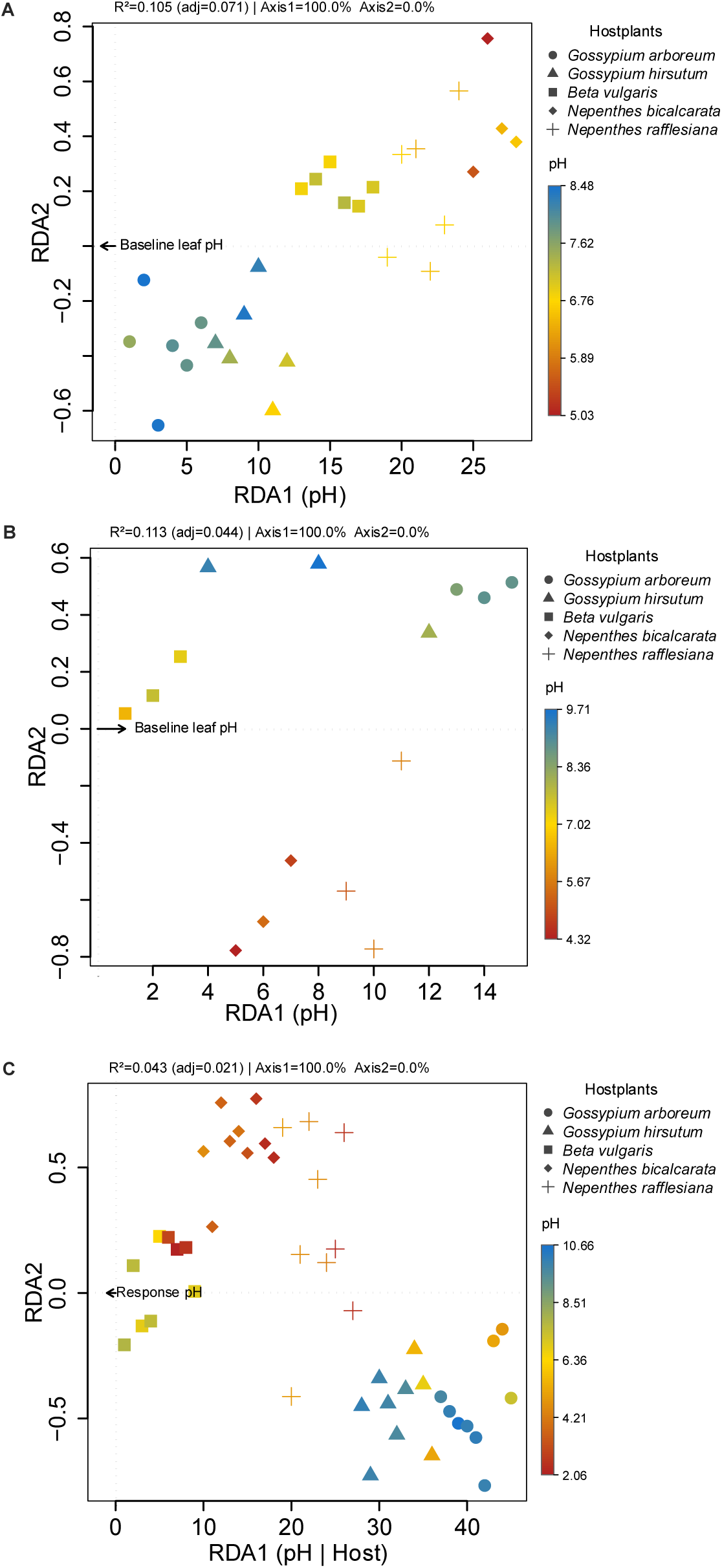
Apparent pH associations in unconditioned models are not supported after accounting for host-structured sampling. RDA models were fit using reaction-level functional profiles constrained by pH without conditioning on host species. Because samples are grouped by host species in the uninoculated leaves (accession GSE281272), significance for Panels B and C was evaluated using permutations restricted within host species (block-stratified permutations). (A) Inoculated leaves (present study). Baseline pH was significantly associated with functional composition in the unconditioned model (F = 3.07, P = 0.005; R² = 0.1055, adj. R² = 0.0711). (B) Uninoculated dry leaves (NCBI GEO accession GSE281272, control dataset). No significant pH association was detected when permutations were restricted within host species (F = 1.65, P = 0.459; R² = 0.1126, adj. R² = 0.0443). (C) Uninoculated leaves exposed to pH spray treatments (NCBI GEO accession GSE281272). No significant pH association was detected when permutations were restricted within host species (F = 1.94, P = 0.68; R² = 0.0433, adj. R² = 0.0210).

**Supplementary Figure S2.**
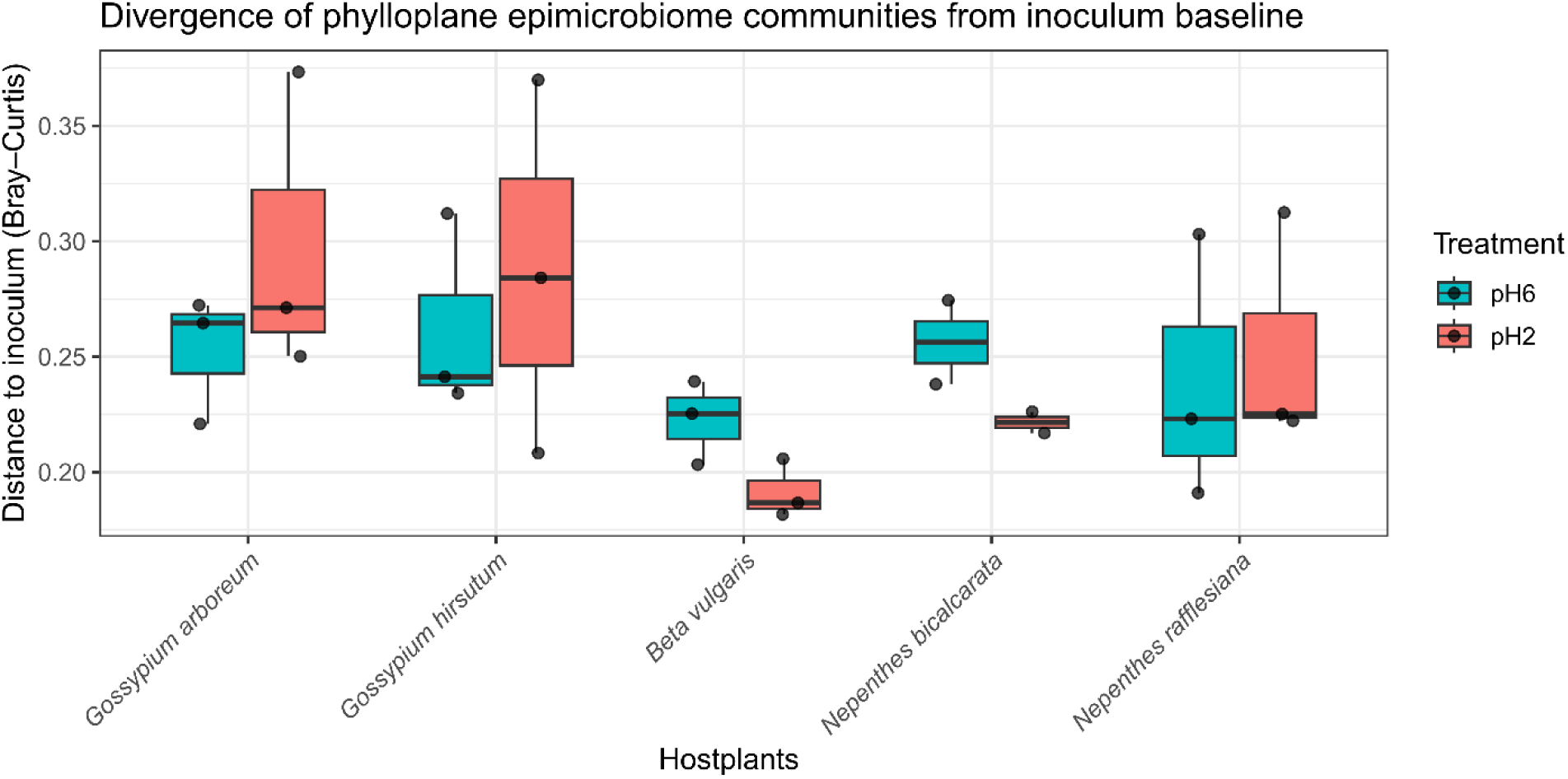
Divergence of phylloplane epimicrobiome communities from inoculum baseline across hostplants and pH treatment. The Bray-Curtis dissimilarity between phylloplane microbial communities and the soil inoculum was calculated for each hostplant under two pH conditions (pH6 and pH2) Boxplots represent the distribution of distances across biological replicates, where the center line indicates the median. Points represent individual samples.

**Supplementary Figure S3.**
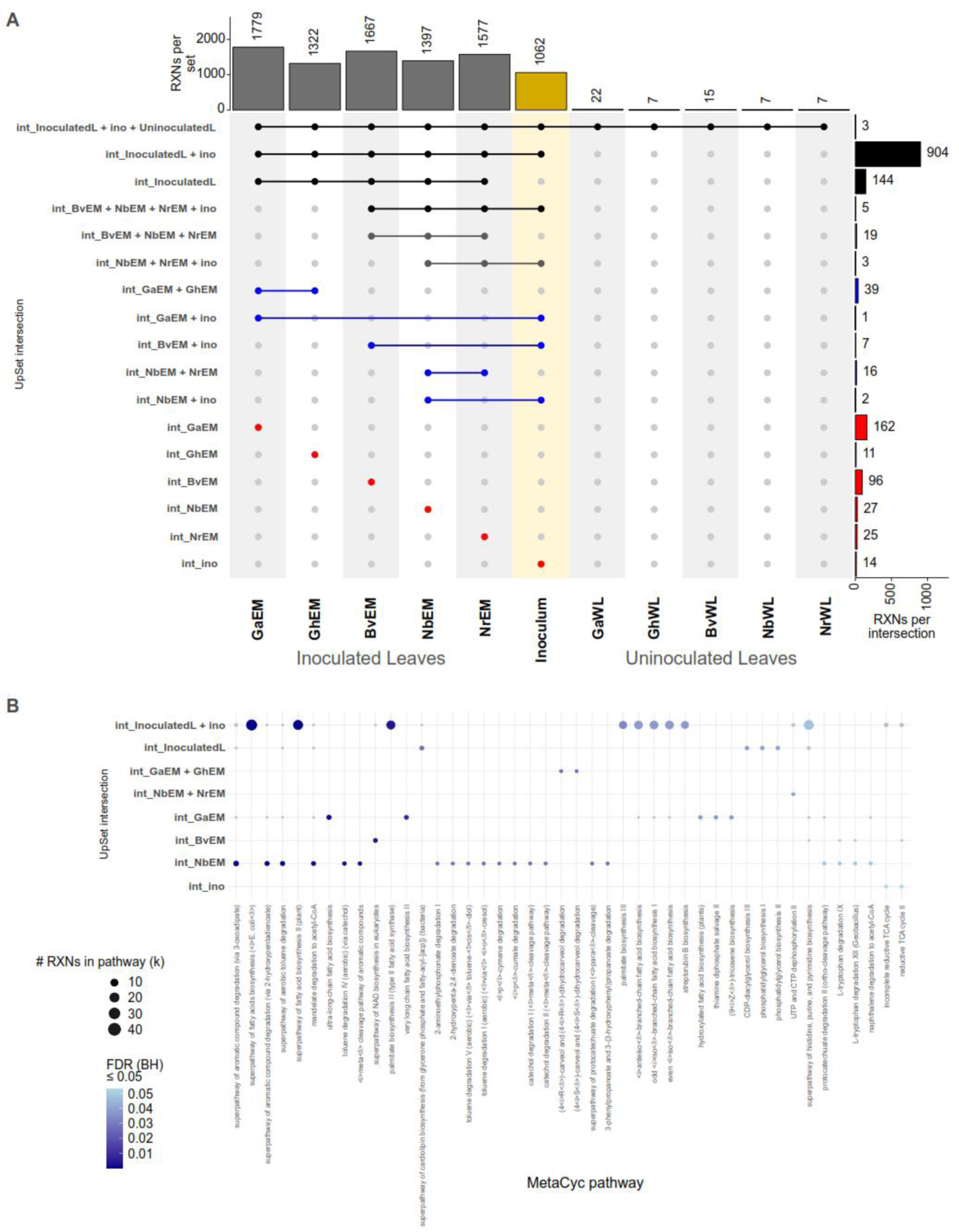
Reaction intersections and pathway enrichment under stricter filtering criteria. **(A)** UpSet plot of HUMAnN3-derived reactions across inoculated leaves, uninoculated whole-leaf datasets, and the inoculum, using a more conservative presence definition (e.g., increased prevalence threshold and/or higher CPM cutoff). Top bars indicate total reactions per group (set size), and right bars indicate intersection size. **(B)** MetaCyc pathway enrichment analysis for intersections containing ≥10 reactions under the stricter filtering scheme. Enrichment was tested using a one-sided hypergeometric test with the background defined as the union of annotated reactions across tested intersections; P-values were adjusted using the Benjamini-Hochberg method. Only pathways with FDR-adjusted q ≤ 0.05 are shown. This analysis demonstrates robustness of intersection structure and functional enrichment patterns to parameter stringency. Major patterns observed in Figure 4 were preserved under conservative filtering, indicating that conclusions are not driven by low-prevalence reactions.

**Supplementary Figure S4.**
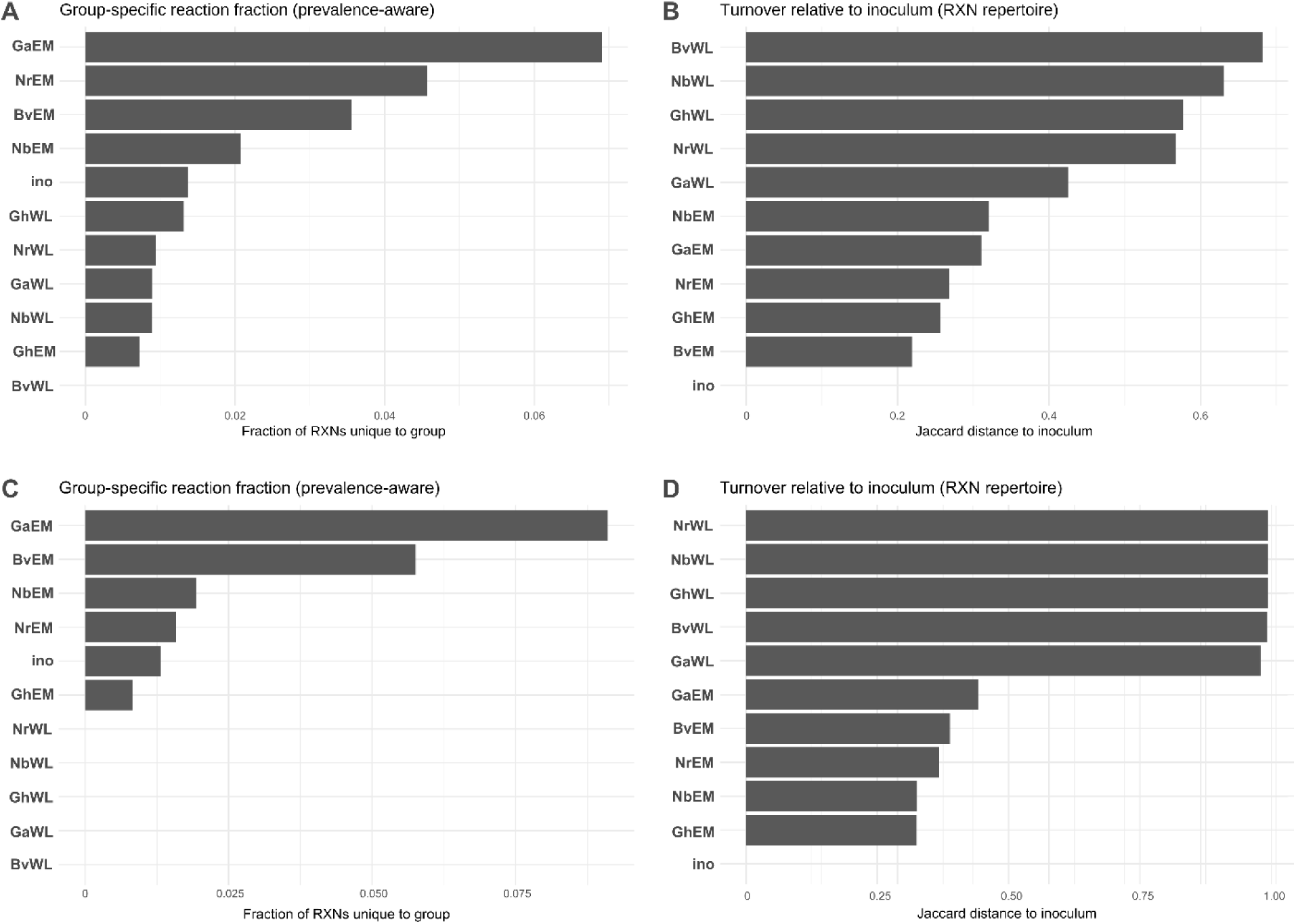
Prevalence-aware estimation of group-specific reaction fractions and functional turnover of leaf communities relative to the soil inoculum. (A) Fraction of reactions unique to each group (GaEM, GhEM, BvEM, NbEM, NrEM, ino, GaWL, GhWL, BvWL, NbWL, NrWL) after applying prevalence-aware presence filtering (reaction considered present if CPM ≥ 1 in ≥66% of samples within a group). Reactions were derived from HUMAnN3 CPM-normalized outputs and converted to binary presence–absence matrices prior to intersection analysis. Bars represent the proportion of reactions detected exclusively in a given group relative to the total reactions observed across all groups. This analysis quantifies functional uniqueness independent of abundance magnitude and complements the intersection-based visualization in Figure 4A. (C) As above, but with stricter filters of Supplementary Figure S3. (B) Jaccard distance of reaction repertoires between each group and the soil inoculum (ino), calculated from prevalence-filtered presence–absence matrices. Values range from 0 (identical reaction sets) to 1 (completely distinct repertoires). Higher values indicate greater functional turnover relative to the inoculum. This analysis evaluates the extent to which leaf-associated microbiomes retain or diverge from inoculum-derived functional potential. (D) As above, but with stricter filters of Supplementary Figure S3.

**Supplementary Figure S5.**
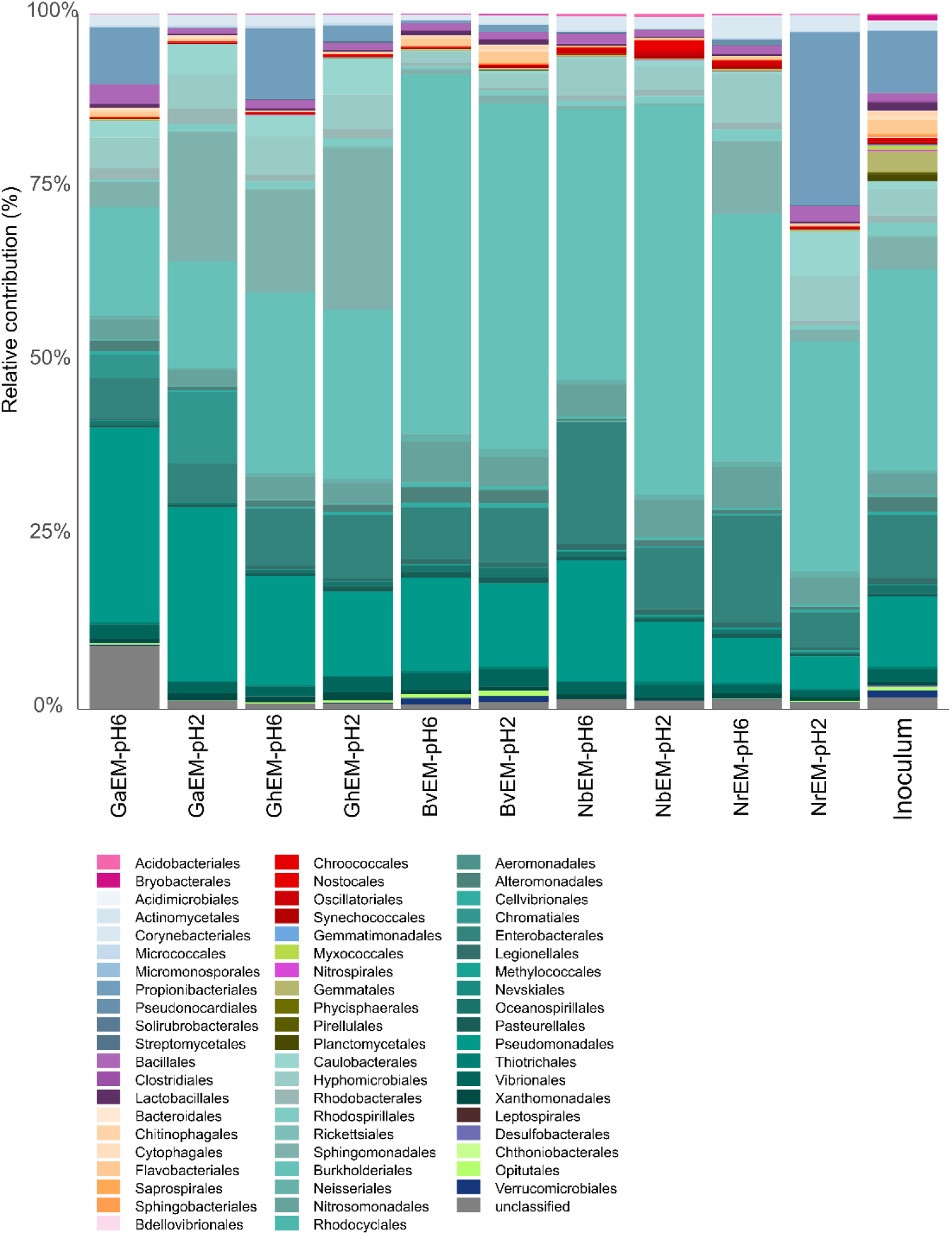
Order-level contribution to total reaction abundance across hostplant-treatment groups and inoculum. Stacked bar plots show the relative contribution (%) of microbial taxonomic orders to the full HUMAnN3 reaction abundance table (reaction abundances stratified by taxonomy and collapsed to order level). For each hostplant epimicrobiome (EM)-treatment group (GaEM-pH6, GaEM-pH2, GhEM-pH6, GhEM-pH2, BvEM-pH6, BvEM-pH2, NbEM-pH6, NbEM-pH2, NrEM-pH6, NrEM-pH2) and inoculum (ino), reaction abundances were summed across all reactions assigned to each taxonomic order. Order-level totals were then converted to percentages such that contributions sum to 100% within each group. “Unclassified” denotes reaction abundances not resolved to a specific taxonomic order in the stratified HUMAnN3 output.

**Supplementary Dataset S1. Differentially abundance features comparing epimicrobiome leaves against inoculum**

## Notes

### Competing Interest Statement

The authors have declared no competing interest.

